# pVACtools: a computational toolkit to identify and visualize cancer neoantigens

**DOI:** 10.1101/501817

**Authors:** Jasreet Hundal, Susanna Kiwala, Joshua McMichael, Christopher A. Miller, Alexander T. Wollam, Huiming Xia, Connor J. Liu, Sidi Zhao, Yang-Yang Feng, Aaron P. Graubert, Amber Z. Wollam, Jonas Neichin, Megan Neveau, Jason Walker, William E Gillanders, Elaine R. Mardis, Obi L. Griffith, Malachi Griffith

## Abstract

Identification of neoantigens is a critical step in predicting response to checkpoint blockade therapy and design of personalized cancer vaccines. We have developed an *in silico* sequence analysis toolkit - pVACtools, to facilitate comprehensive neoantigen characterization. pVACtools supports a modular workflow consisting of tools for neoantigen prediction from somatic alterations (pVACseq and pVACfuse), prioritization and selection using a graphical web-based interface (pVACviz) and design of DNA vector-based vaccines (pVACvector) and synthetic long peptide vaccines. pVACtools is available at pvactools.org.

---

The increasing use of cancer immunotherapies has spurred interest in identifying and characterizing predicted neoantigens encoded by a tumor genome. The facility and precision of computational tools for predicting neoantigens have become increasingly important(1), and several such resources have been published(2–4). Typically, these tools start with a list of somatic variants (in VCF or other formats) with annotated protein changes, and predict the strongest MHC binding peptides (8-11-mer for class I MHC and 13-25-mer for class II) using one or more prediction algorithms(5–7). The predicted neoantigens are then filtered and ranked based on defined metrics including sequencing read coverage, variant allele fraction (VAF), gene expression, and differential binding compared to the wild type peptide (agretopicity index score(8)). However, of the small number of such prediction tools **(Supp Table 1)**, most lack key functionality, including predicting neoantigens from gene fusions, aiding optimized vaccine design for DNA cassette vaccines, and incorporating nearby germline or somatic alterations into the candidate neoantigens(9). Furthermore, none of the existing tools offer an intuitive graphical user interface for visualizing and efficiently selecting the most promising candidates; a key feature for facilitating involvement of clinicians and other researchers in the process of neoantigen evaluation.

To address these limitations, we created a comprehensive and extensible toolkit for computational identification, selection, prioritization and visualization of neoantigens - *‘*pVACtools*’*, that facilitates each of the major components of neoantigen identification. This computational framework can be used to identify neoantigens from a variety of somatic alterations, including gene fusions and insertion/deletion frameshift mutations, both of which potentially create strong immunogenic neoantigens(10). pVACtools can facilitate both MHC class I and II predictions, and provides an interactive display of predicted neoantigens for review by the end user.

The pVACtools workflow (**Figure 1**) is divided into modular components that can be run independently. The main tools in the workflow are: (a) pVACseq: a significantly enhanced and reengineered version of our previous pipeline(11) for identifying and prioritizing neoantigens from a variety of tumor-specific alterations (b) pVACfuse: a tool for detecting neoantigens resulting from gene fusions (c) pVACviz: a graphical user interface web client for process management, visualization and selection of results from pVACseq (d) pVACvector: a tool for optimizing design of neoantigens and nucleotide spacers in a DNA vector that prevents high-affinity junctional neoantigens, and (e) pVACapi: an OpenAPI HTTP REST interface to the pVACtools suite.

**Figure 1:**
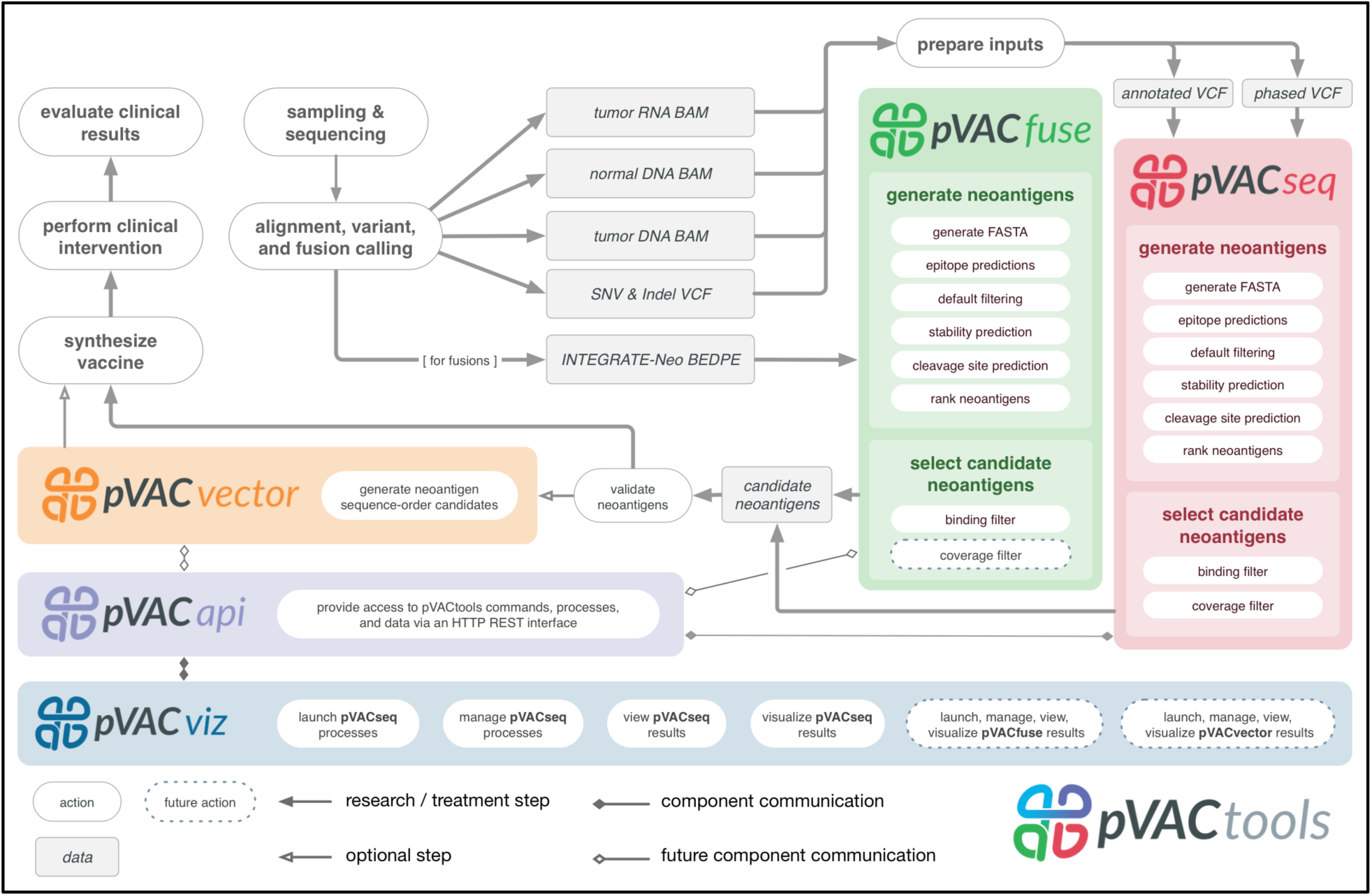
Overview of pVACtools workflow: The pVACtools workflow is highly modularized and is divided into flexible components that can be run independently. The main tools under the workflow include pVACseq for identifying and prioritizing neoantigens from a variety of somatic alterations (red inset box), pVACfuse (green) for detecting neoantigens resulting from gene fusions, pVACviz (blue) for process management, visualization and selection of results and pVACvector (orange) for optimizing design of neoantigens and nucleotide spacers in a DNA vector. All of these tools interact via the pVACapi (purple), an OpenAPI HTTP REST interface to the pVACtools suite.

pVACseq(11) has been re-implemented in Python3 and extended to include many new features since our initial report of its use. pVACseq no longer requires a custom input format for variants, and now uses a standard VCF file annotated with VEP(12). In our own neoantigen identification pipeline, this VCF is the result of merging results from multiple somatic variant callers and RNA expression tools (**Supplementary Text/Example pipeline for creation of pVACtools input files**). Information that is not natively available in the VCF output from somatic variant callers (such as coverage and variant allele fractions for RNA and DNA, as well as gene and transcript expression values) can be added to the VCF using VAtools (http://vatools.org), a suite of accessory routines that we created to accompany pVACtools. pVACtools queries these features directly from the VCF, enabling prioritization and filtering of neoantigen candidates based on sequence coverage and expression information. In addition, pVACseq now makes use of phasing information taking into account variants proximal to somatic variants of interest. Since proximal variants can change the peptide sequence and affect neoantigen binding predictions, this is important for ensuring that the selected neoantigens correctly represent the individual’s genome (9). We have also expanded the supported mutation types for neoantigen predictions to include in-frame indels and frameshift mutations. These capabilities expand the potential number of targetable neoantigens several-fold in many tumors (10,13)(**Supplementary Data**).

To prioritize neoantigens, pVACseq now offers support for eight different MHC Class I antigen prediction algorithms and four MHC Class II prediction algorithms. The tool does this in part by leveraging the Immune Epitope Database (IEDB)(14) and their suite of six different MHC class I prediction algorithms, as well as three MHC Class II algorithms (**Methods/Neoantigen prediction**). pVACseq supports local installation of these tools for high-throughput users, access through a docker container (https://hub.docker.com/r/griffithlab/pvactools), or provides ready-to-go access via the IEDB RESTful web interface. In addition, pVACseq now contains an extensible framework for supporting new neoantigen prediction algorithms that has been used to add support for two new non-IEDB algorithms - MHCflurry(15) and MHCnuggets(16). By creating a framework that integrates many tools we allow for (a) a broader ensemble approach than IEDB, and (b) a system that other users can leverage to develop improved ensemble ranking, or to integrate proprietary or not-yet-public prediction software. Importantly, this framework enables non-informatics-expert users to predict neoantigens from sequence variant data sets.

Once neoantigens have been predicted, the pVACseq ranking score is used to prioritize them. This score takes into account gene expression, sequence read coverage, binding affinity predictions, and agretopicity (**Methods/Ranking of Neoantigens**). In addition to applying strict binding affinity cutoffs, the pipeline also offers support for MHC allele-specific cutoffs(17). We also offer cleavage position predictions via optional processing through NetChop(18) as well as stability predictions made by NetMHCstabPan(19).

Previous studies have shown that the novel protein sequences produced by gene fusions frequently produce neoantigen candidates(20). pVACfuse provides support for predicting neoantigens from such gene fusions. Fusion variants may be imported in annotated BEDPE format from any fusion caller. We recommend using INTEGRATE-Neo(20) for annotation of fusion calls in BEDPE format. These variants are then assessed for presence of fusion neoantigens using predictions from any of the pVACseq-supported binding prediction algorithms.

Implementing cancer vaccines in a clinical setting requires multidisciplinary teams, many of whom may not be informatics experts. To support this growing community of users, we developed pVACviz, which is a browser-based user interface that assists in launching, managing, reviewing, and visualizing the results of pVACtools processes. Instead of interacting with the tools via terminal/shell commands, the pVACviz client provides a modern web-based user experience. Users complete a pVACseq (**Figure 2**) process setup form that provides helpful documentation and suggests valid values for inputs. The client also provides views showing ongoing processes, their logs, and interim data files to aid in managing and troubleshooting. After a process has completed, users may examine the results as a filtered data table, or as a scatterplot visualization - allowing them to curate results and save them as a CSV file for further analysis. Extensive documentation of the visualization interface can be found in the online documentation (https://pvactools.readthedocs.io/en/latest/pvacviz.html).

**Figure 2:**
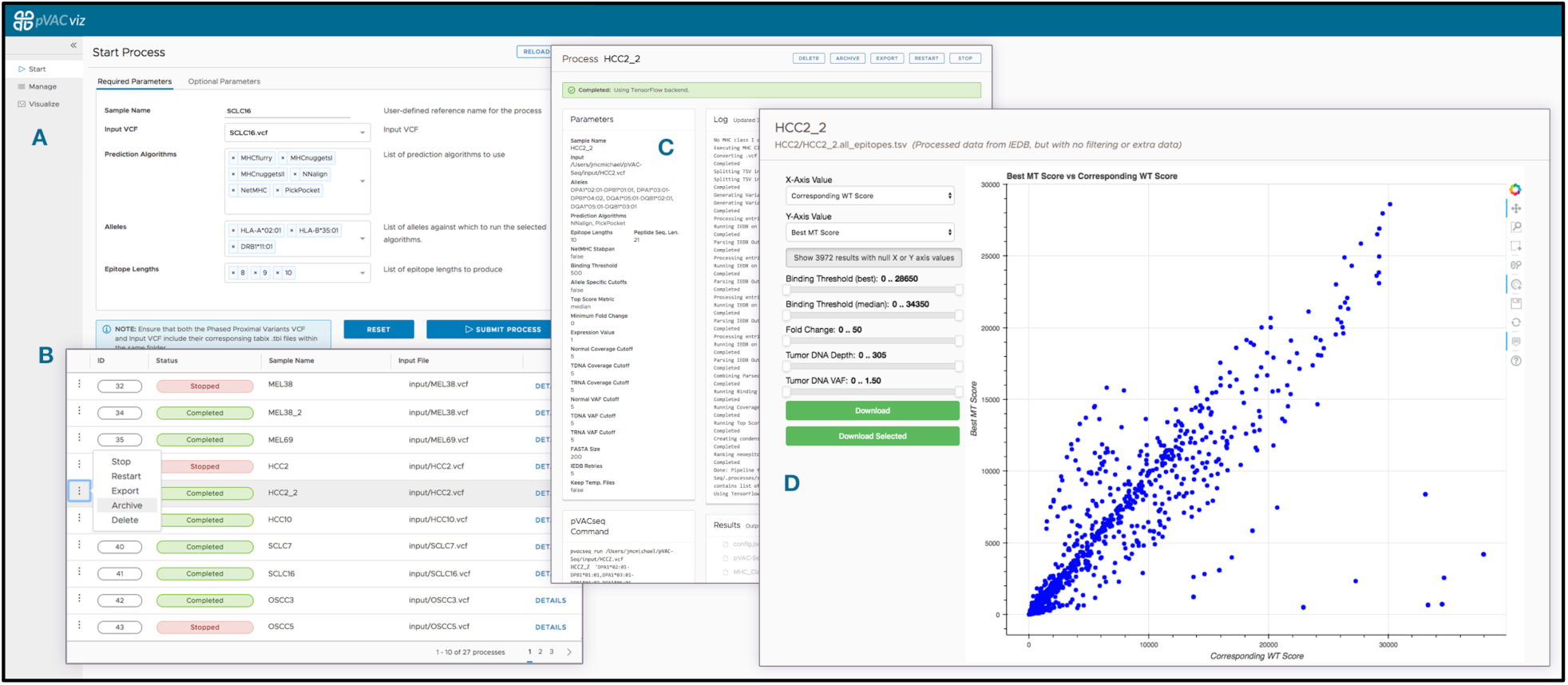
pVACviz GUI client: pVACtools provides a browser-based graphical user interface, called pVACviz, that provides an intuitive means to launch pipeline processes, monitor their execution, and analyze, export, or archive their results. To launch a process, users navigate to the Start Page (A), and complete a form containing all of the relevant inputs and settings for a pVACseq process. Each form field includes help text, and provides typeahead completion where applicable. For instance, the Alleles field provides a typeahead dropdown menu that matches available alleles. Once a process is launched, a user may monitor its progress on the Manage Page (B), which lists all running, stopped, and completed processes. The Details Page (C) shows a process’ current log, attributes, and any results files as well as providing controls for stopping, restarting, exporting and archiving the process. The results of pipeline processes may be analyzed on the Visualize Page (D), which displays a customizable scatterplot of a file’s rows. The X and Y axis may be set to any column in the result set, and filters may be applied to values in any column. Additionally, points may be selected on the scatter plot or data grid (not visible in this figure) for further analysis or export as CSV files.

Furthermore, to support informatics groups that want to incorporate or build upon the pVACtools features, we developed pVACapi, which provides a HTTP REST interface to the pVACtools suite. Currently, it provides the API that pVACviz uses to interact with the pVACtools suite. Advanced users could develop their own user interfaces, or use the API to control multiple pVACtools installations remotely over an HTTP network.

Once a list of neoantigen candidates has been prioritized and selected, the pVACvector utility can be used to aid in the construction of DNA-based cancer vaccines. The input is either the output file from pVACseq or a fasta file containing peptide sequences. pVACvector returns a neoantigen sequence ordering that minimizes the effects of junctional peptides (which may create novel antigens) between the sequences (**Figure 3)**. This is accomplished by using the core pVACseq module to predict the binding scores for each junctional peptide and by modifying junctions with spacer(21,22) amino acid sequences, or by trimming the ends of the peptides in order to reduce reactivity. The final vaccine ordering is achieved through a simulated annealing procedure that returns a near-optimal solution, when one exists (**Methods/Implementation**).

**Figure 3:**
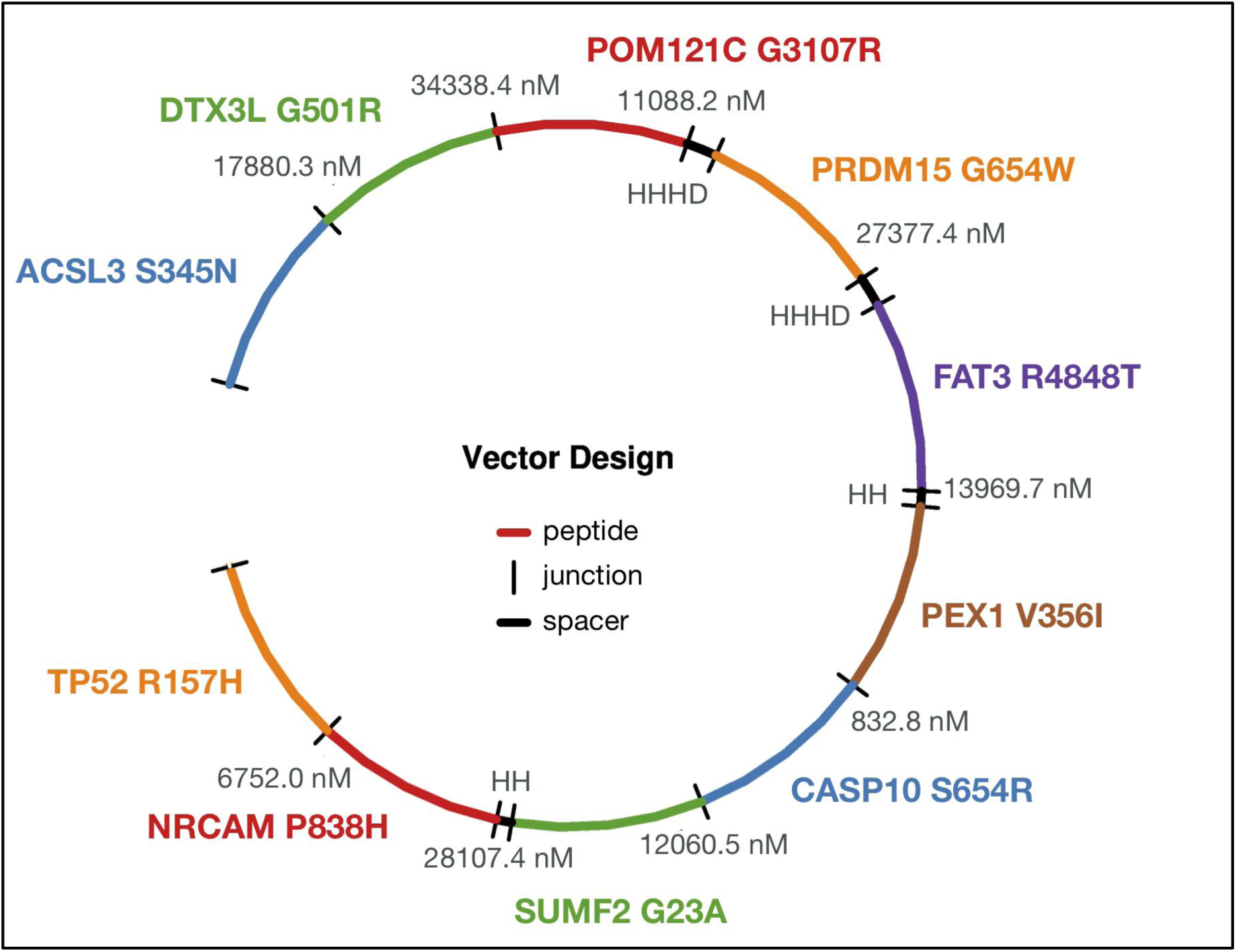
An example pVACvector output showing the optimum arrangement of candidate neoantigens for a DNA-vector based vaccine design. The figure depicts a circularized DNA insert carrying 10 encoded neoantigenic peptide sequences to be synthesized and encoded/cloned into a DNA plasmid. DNA sequences encoding each peptide are ordered (with use of spacer sequences where needed) to ensure there are no strong-binding junctional epitopes. Each neoantigenic peptide candidate is shown in Blue, Green, Red, Orange, Purple, and Brown. Spacer sequences, where added to minimize junctional epitope affinity, are depicted in Black, along with the binding affinity value of the junctional epitope. Labels represent the Gene Name and Amino Acid Change for each candidate.

As many prediction algorithms are CPU-intensive, pVACseq, pVACfuse, and pVACvector also support using multiple cores to improve runtime. Using this feature, calls to IEDB and other prediction algorithms are made in parallel over a user-defined number of processes. (**Methods/Implementation**).

pVACtools has been used to predict and prioritize neoantigens for several immunotherapy studies(23–25) and cancer vaccine clinical trials (e.g. NCT02348320, NCT03121677, NCT03122106, NCT03092453 and NCT03532217). We also have a large external user community that has been actively evaluating and using these packages for their neoantigen analysis, and has also helped in the subsequent refinement of pVACtools through extensive feedback. The original ‘pvacseq’ package has been downloaded over 41,000 times from PyPi, and the ‘pvactools’ package has been downloaded over 18,000 times.

To demonstrate the utility and performance of the pVACtools package, we downloaded exome sequencing and RNA-Seq data from The Cancer Genome Atlas (TCGA)(26) from 100 cases each of melanoma, hepatocellular carcinoma and lung squamous cell carcinoma, and used patient-specific MHC Class I alleles (**Supp Fig 1**) to determine neoantigen candidates for each tumor. By extending support for additional variant types (**Supp Fig 2)** as well as prediction algorithms, we produced 42% more predicted neoantigens compared to the previous version of pVACseq(11) **(Supplementary Text/Analysis of TCGA data using pVACtools)**.

## Comparison of epitope prediction software

Since we offer support for as many as eight different epitope prediction tools, we assessed agreement in binding affinity predictions (IC50) between these algorithms from a random subset of 100,000 neoantigen peptides from the TCGA analysis (**Figure 4**). The highest correlation was observed between the two stabilization matrix method (SMM)-based algorithms - SMM and SMMPMBEC. The next best correlation was observed between NetMHC and MHCflurry, possibly due to both being allele-specific predictors employing neural network based models. Overall the correlation between prediction algorithms is low (mean correlation of 0.388 and range of 0.18-0.89 between all pairwise comparisons of algorithms).

**Figure 4:**
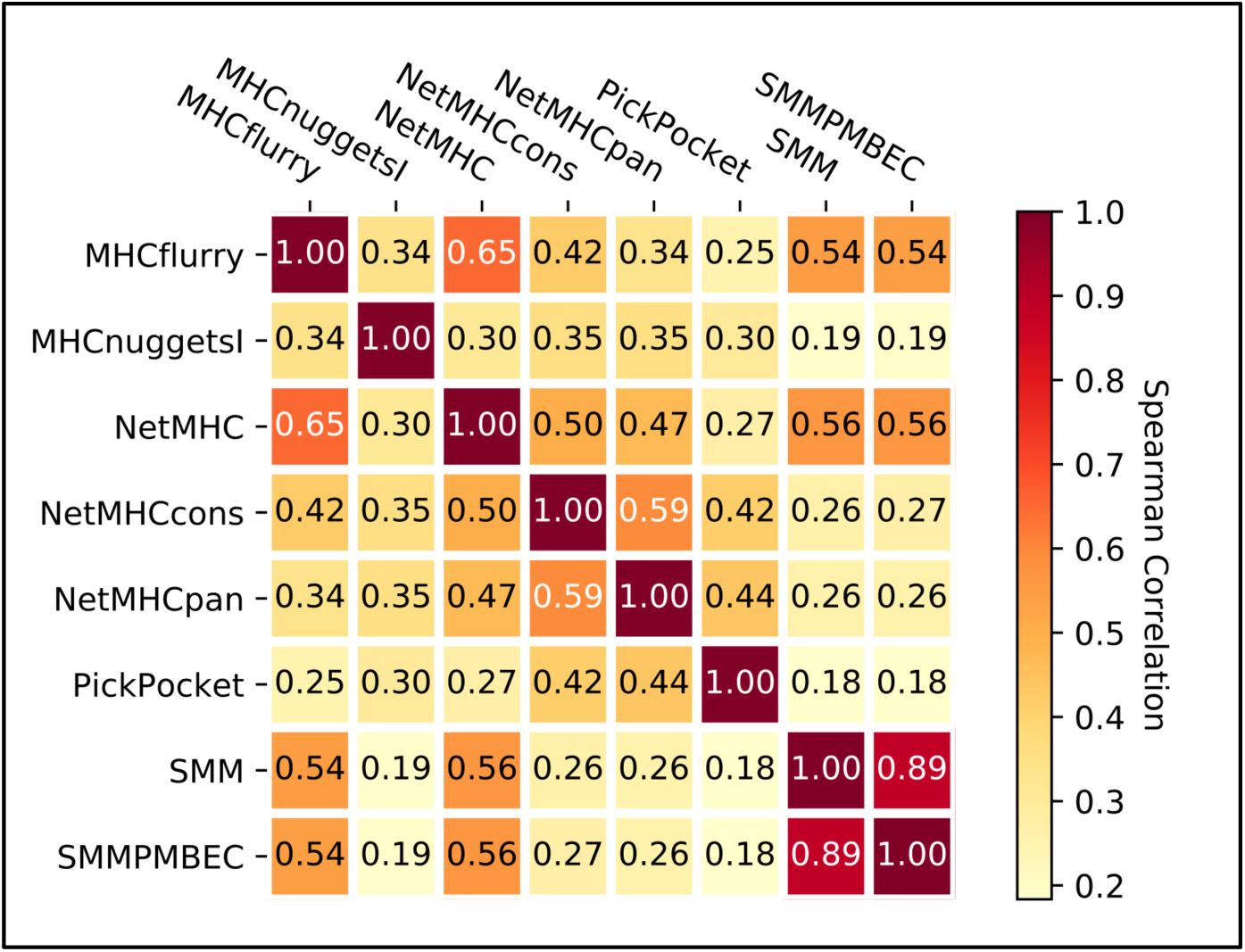
Correlation between prediction algorithms: Spearman Correlation between prediction values from all 8 class I prediction algorithms generated from a random subsample of 100,000 peptides.

We also evaluated if there were any biases among the algorithms to predict strong (i.e. binding affinity < = 500nM) or weak binding epitopes (**Figure 5 and Supp Fig 3**). We found that MHCnuggets predicts the highest number of strong-binding candidates alone. Of the total number of strong binding candidates predicted, 64.7% of these candidates were predicted by a single algorithm (any one of the eight algorithms), 35.2% were predicted as strong-binders by two to seven algorithms, and only 1.8% of the strong-binding candidates were predicted as strong binders by the combination of all eight algorithms. In fact, as shown in **Figure 6**, even if one (or more) algorithms predict a peptide to be a strong binder, often another algorithm not only doesn’t agree but disagrees by a large margin, in some cases predicting that same peptide as a very weak binder. This remarkable lack of agreement underscores the potential value of an ensemble approach that considers multiple algorithms.

**Figure 5:**
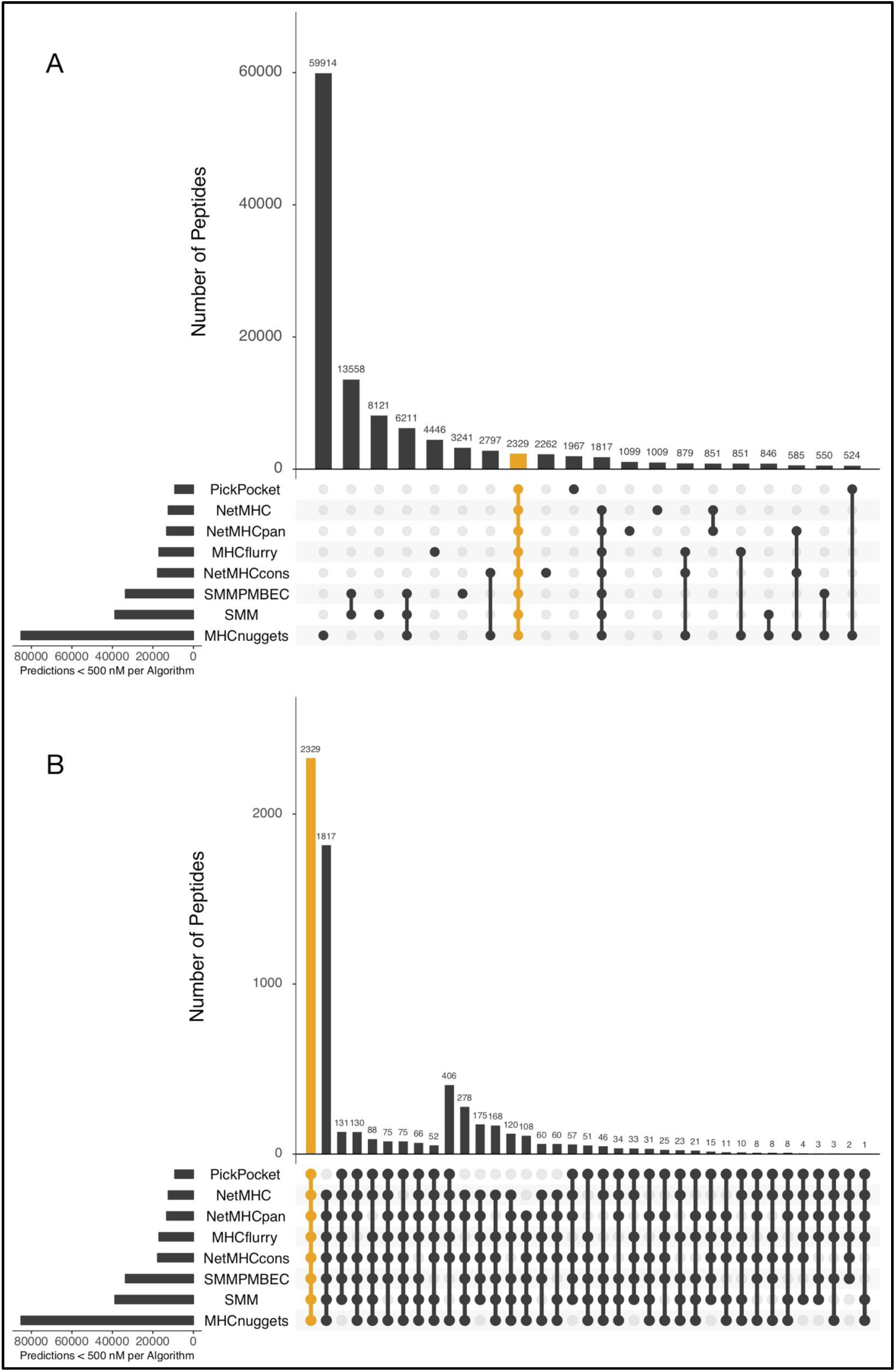
Intersection of peptide sequences predicted by different algorithms are shown using upset plots. The y-axis shows the number of overlapping unique neoantigenic peptides predicted for each combination of algorithm depicted on the x-axis. Each collection of connected circles shows the set contained in an exclusive intersection (i.e. the identity of each algorithm), while the light gray circles represent the algorithm(s) that do not participate in this exclusive intersection. (a) Upset plot for the top 20 algorithm combinations ranked by the number of peptides predicted to be a good binder (mutant IC50 score < 500 nM). The combination of all eight algorithms (highlighted orange) ranks the 8th highest; (b) Upset plot for algorithm combinations where at least six algorithms agree on predicting a peptide to be a good binder (MT IC50 score < 500 nM). The combination of all eight algorithms (highlighted orange) ranks the highest.

**Figure 6:**
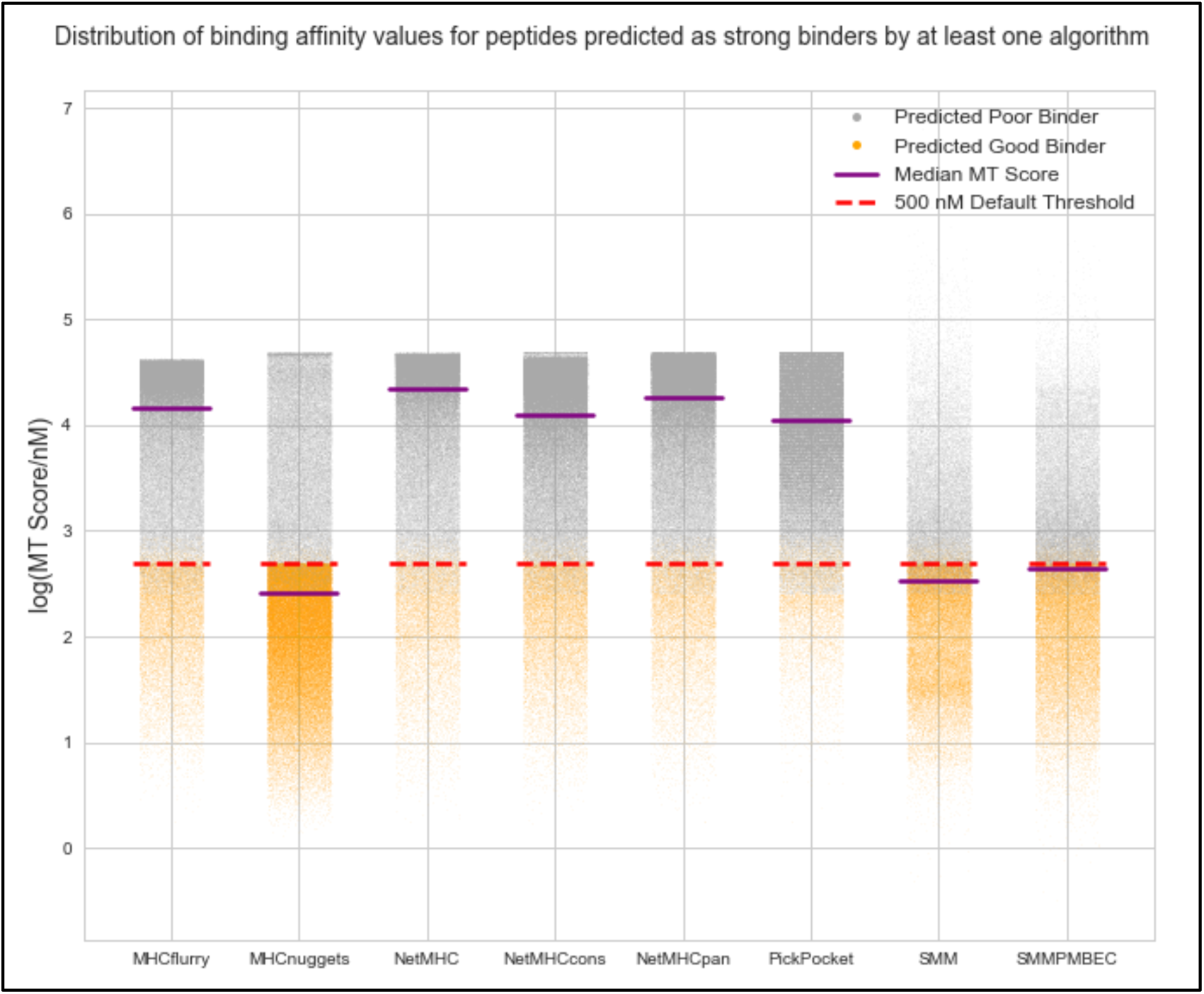
Overall distribution of binding affinity scores (nM) for 126,648 peptides (out of total predicted 14,599,993 peptides) where at least one of the algorithms predicts a strong binder. To define the set of peptides that are strong binders according to at least one algorithm, HLA allele subtype-specific thresholds were applied when available, otherwise the default cutoff binding affinity of 500 nM was used. The peptides were further filtered using the default coverage based filters. Peptides with predicted MT IC50 scores lower than their respective cutoff scores are highlighted in orange. The median MT IC50 scores of each algorithm’s prediction are indicated for reference (purple line).

Next we determined if the number of human HLA alleles supported by these eight algorithms differed. As shown (**Supp Fig 4)**, MHCnuggets supports the highest number of human HLA alleles.

As reported from our demonstration analysis, a typical tumor has too many possible neoantigen candidates to be practical for a vaccine. There is therefore a critical need for a tool that takes in the input from a tumor sequencing analysis pipeline and reports a filtered and prioritized list of neoantigens. pVACtools enables a streamlined, accurate and user-friendly analysis of neoantigenic peptides from NGS cancer datasets. This suite offers a complete and easily configurable end-to-end analysis, starting from somatic variants and gene fusions (pVACseq and pVACfuse respectively), through filtering, prioritization, and visualization of candidates (pVACviz), and determining the best arrangement of candidates for a DNA vector vaccine (pVACvector). Furthermore, by supporting additional classes of variants as well as gene fusions, we offer an increase in the number of predicted neoantigens which may be critical for low mutational burden tumors. Finally, by extending support for multiple binding prediction algorithms, we allow for a consensus approach. The need for this integrated approach is made abundantly clear by the high disagreement between these algorithms observed in our demonstration analyses.

The results from pVACtools analyses are already being used in dozens of cancer immunology studies, including studying the relationship between tumor mutation burden and neoantigen load to predict response in checkpoint blockade therapy trials and the design of cancer vaccines in ongoing clinical trials. We anticipate that pVACtools will make such analyses more robust, reproducible, and facile as these efforts continue.

## Methods

### TCGA data pre-processing

Aligned (build GRCh38) tumor and normal BAMs from BWA(27) (version 0.7.12-r1039) as well as somatic variant calls from VarScan2(28,29)(in VCF format) were downloaded from the Genomic Data Commons (GDC, https://gdc.cancer.gov/). Since the GDC does not provide germline variant calls for TCGA data, we used GATK’s(30) HaplotypeCaller to perform germline variant calling using default parameters. These calls were refined using VariantRecalibrator in accordance with GATK Best Practices(31). Somatic and germline missense variant calls from each sample were then combined using GATK’s CombineVariants, and the variants were subsequently phased using GATK’s ReadBackedPhasing algorithm.

Phased Somatic VCF files were annotated with RNA depth and expression information using VAtools (http://vatools.org). We restricted our analysis to only consider ‘PASS’ variants in these VCFs as these are higher confidence than the raw set, and the variants were annotated using the “--pick” option in VEP (Ensembl version 88).

Existing *in silico* HLA typing information was obtained from The Cancer Immunome Atlas (TCIA) database(32).

### Neoantigen prediction

The VEP-annotated VCF files were then analyzed with pVACseq using all eight Class I prediction algorithms for neoantigen peptide lengths 8-11. The current MHC Class I algorithms supported by pVACseq are NetMHCpan(33), NetMHC(7,33), NetMHCcons(34), PickPocket(35), SMM(36), SMMPMBEC(37), MHCflurry(15) and MHCnuggets(16). The four MHC Class II algorithms that are supported are NetMHCIIpan, SMMalign, NNalign, and MHCnuggets. For the demonstration analysis, we limited our prediction to only MHC Class I alleles due to availability of HLA typing information from TCIA, though binding predictions for Class II alleles can also be generated using pVACtools.

### Ranking of Neoantigens

To help prioritize neoantigens, a ranking score is assigned to all neoantigens that pass initial filters where each of the following four criteria are assigned a rank-ordered value (where the worst = 1):

B = binding affinity

F = Fold Change between MT and WT alleles

M = mutant allele expression, calculated as (Gene expression * Mutant allele RNA Variant allele fraction)

D = DNA Variant allele fraction

A final ranking is based on a score obtained from combining these values: Priority Score = B+F+(M*2)+(D/2). This score is not meant to be a definitive metric of peptide suitability for vaccines, but was designed to be a useful first step in the peptide selection process.. Moreover, since the score is based on rank-ordered values, each neoantigen’s score is relative to the scores of the other neoantigens it is evaluated against and can not be used to compare neoantigens between different pVACseq runs.

### Implementation

pVACtools is written in Python3. The individual tools are implemented as separate command line entry points that can be run using the ‘`pvacseq`’, ‘`pvacfuse`’, ‘`pvacvector`’, ‘`pvacapi`’, and ‘`pvacviz`’ commands to run the respective tool. pVACapi is required to run pVACviz so both the ‘pvacapi’ and ‘pvacviz’ commands need to be executed in separate terminals.

The test suite is implemented using the Python unittest framework and GitHub integration tests are run using travis-ci (travis-ci.org). Code changes are integrated using GitHub pull requests (https://github.com/griffithlab/pVACtools/pulls). Feature additions, user requests, and bug reports are managed using the GitHub issue tracking (https://github.com/griffithlab/pVACtools/issues). User documentation is written using the reStructuredText markup language and the Sphinx documentation framework (sphinx-doc.org). Documentation is hosted on Read The Docs (readthedocs.org). The `pymp-pypi` package was used to add support for parallel processing. The number of processes is controlled by the `--n-threads` parameter.

#### VACseq

For pVACseq, the `pyvcf` package is first used for parsing the input VCF file and extracting information about the supported missense, inframe indel, and frameshift mutations into TSV format.

This output is then used to determine the wildtype peptide sequence by extracting a region around the somatic mutation according to the `--peptide-sequence-lengt`h specified by the user. The mutation’s amino acid change is incorporated in this peptide sequence to determine the mutant peptide sequence. For frameshift mutations, the new downstream protein sequence calculated by VEP is reported from the mutation position onward. The number of downstream amino acids to include is controlled by the `--downstream-sequence-length` parameter. If a phased VCF with proximal variants is provided, proximal missense mutations that are in phase with the somatic variant of interest are incorporated into the mutant and wildtype peptide sequences as appropriate. The mutant and wildtype sequences are stored in a FASTA file. The FASTA file is then submitted to the individual prediction algorithms for binding affinity predictions. For algorithms included in IEDB, we either use the IEDB API or a standalone installation, if an installation path is provided by the user (`--iedb-install-directory`). The `mhcflurry` and `mhcnuggets` packages are used to run the MHCflurry(15) and MHCnuggets(16) prediction algorithms, respectively.

The predicted mutant antigens are then parsed into a TSV report format and for each mutant antigen the closest wildtype antigen is determined and reported. Predictions for each mutant antigen/neoantigen from multiple algorithms are aggregated into the “best” (lowest) and median binding scores. The resulting TSV is processed through multiple filtering steps (**Supplementary Text/Comparison of filtering criteria)**: (1) Binding filter: this filter selects the strongest binding candidates based on the mutant binding score and the fold change (WT score / MT score). Depending on the `--top-score-metric` parameter setting, this filter is either applied to the median score across all chosen prediction algorithms(default) or the best score amongst the chosen prediction algorithms. (2) Coverage filter: this filter accepts VAF and coverage information from the tumor DNA, tumor RNA, and normal DNA, if these values are available in the input VCF. (3) Transcript-support-level (TSL) filter: this filter evaluates each transcript’s support level if this information was provided by VEP in the VCF. (4) Top-score filter: the filter picks the top mutant peptide for each variant, using the binding affinity as the determining factor. This filter is implemented to only select the best candidate from amongst multiple candidates that could result from a single variant due to different peptide lengths, variant registers, transcript sequences, and HLA alleles. The result of these filterings steps is reported in a filtered report TSV. The remaining neoantigens are then annotated with cleavage site and stability predictions by NetChop and NetMHCStabPan, respectively, and a relative ranking score (**Methods**/**Ranking of Neoantigens**) is assigned. The rank ordered final output is reported in a condensed file. The `pandas` package is used for data management while filtering and ranking the neoantigen candidates.

#### pVACvector

When running pVACvector with a pVACseq output file, the original input VCF must also be provided (`--input-vcf` parameter). The VCF is used to extract a larger peptide sequence around the target neoantigen (length determined by the `--input-n-mer` parameter). Alternatively, a list of target peptide sequences can be provided in a fasta file. The set of peptide sequences are then combined in all possible pairs, and a ordering of peptides for the vector is produced as follows:

To determine the optimal order of peptide-spacer-peptide combinations, binding predictions are made for all peptide registers overlapping the junction. A directed graph is then constructed, with nodes defined as target peptides, and edges representing junctions. The score of each edge is defined as the lowest binding score of its junctional peptides (a conservative metric). Edges with scores below the threshold are removed, and if heuristics indicate that a valid graph may exist, a simulated annealing procedure is used to identify a path through the nodes that maximizes junction scores (preserving the weakest overall predicted binding for junctional epitopes). If no valid ordering is found, additional “spacer” amino acids are added to each junction, binding affinities are re-calculated, and a new graph is constructed and tested, setting edge weights equal to that of the best performing (highest binding score) peptide-junction-spacer combination.

The spacers used for pVACvector are set by the user with the `--spacers` parameter. This parameter defaults to `None,AAY,HHHH,GGS,GPGPG,HHAA,AAL,HH,HHC,HHH,HHHD,HHL,HHHC`, where `None` is the placeholder for testing junctions without a spacer sequence. Spacers are tested iteratively, starting with the first spacer in the list, and adding subsequent spacers if no valid path is found.

If no result is found after testing with the full set of spacers, the ends of “problematic” peptides, where all junctions contain at least one well-binding epitope, will be clipped by removing one amino acid at a time, then repeating the above binding and graph-building process. This clipping may be repeated up to the number of times specified in the `-- max-clip-length` parameter.

#### pVACapi and pVACviz

pVACapi is implemented using the Python libraries Flask and Bokeh. The pVACviz client is written in TypeScript using the Angular web application framework, the Clarity UI component library, and the ngrx library for managing application state.

## Data availability

Data from 100 cases each of melanoma, hepatocellular carcinoma and lung squamous cell carcinoma were obtained from TCGA and downloaded via the Genomics Data Commons (GDC). This data can be accessed under dbGaP study accession phs000178. Data for demonstration and analysis of fusion neoantigens was downloaded from the Github repo for Integrate (https://github.com/ChrisMaherLab/INTEGRATE-Vis/tree/master/example).

## Software availability

The pVACtools codebase is hosted publicly on GitHub at https://github.com/griffithlab/pVACtools and https://github.com/griffithlab/BGA-interface-projects (pVACviz). User documentation is available at pvactools.org. This project is licensed under the Non-Profit Open Software License version 3.0 (NPOSL-3.0, https://opensource.org/licenses/NPOSL-3.0). pVACtools has been packaged and uploaded to PyPi under the “pvactools” package name and can be installed on Linux systems by running the ‘pip install pvactools’ command. Installation requires a Python 3.5 environment which can be emulated by using Conda. Versioned Docker images are available on DockerHub (https://hub.docker.com/r/griffithlab/pvactools/).

## Acknowledgements

We thank the patients and their families for donation of their samples and participation in clinical trials. We also thank our rapidly growing user community for testing the software and providing useful input, critical bug reports, as well as suggestions for improvement and new features. We are grateful to Drs. Robert Schreiber, Gavin Dunn and Beatriz Carreno for their expertise and guidance on foundational work on cancer immunology using neoantigens and suggestions on improving the pipeline. Obi Griffith was supported by the National Cancer Institute (NCI) of the National Institutes of Health (NIH) under Award Numbers U01CA209936, U01CA231844 and U24CA237719. Malachi Griffith was supported by the National Human Genome Research Institute (NHGRI) of the NIH under Award Number R00HG007940 and the V Foundation for Cancer Research under Award Number V2018-007.

## Author contributions

JH and SK were involved in all aspects of this study including designing and developing the methodology, and writing the manuscript, with help from CAM, ERM, OLG and MG. JH analyzed and interpreted data with input from HX, CJL, SZ and Y-YF. SK wrote the software code with help from ATW, APG, AZW, MN and JW. JM developed pVACviz and pVACapi with assistance from APG. JN, MN, and CAM developed pVACvector. WEG provided input about clinical needs for vaccine design to help improve the tool features. OG and MG supervised the project and revised the paper. All authors read and approved the final manuscript.

