## Supplementary Data for "pVACtools: a computational toolkit to identify and visualize cancer neoantigens"

### Table of Contents

**Supplementary Text**
**Supplementary Figures**
**Supplementary Table**
**Supplementary References**

#### Supplementary Text

**Example pipeline for creation of pVACtools input files**

pVACtools is designed to support a standard VCF variant file format and thus, should be compatible with many existing variant calling pipelines. However, as a reference, we provide the following description of our current somatic and expression analysis pipeline (manuscript in preparation) which has been implemented using docker, CWL(1), and Cromwell(2). The pipeline consists of workflows for alignment of exome DNA- and RNA-Seq data, somatic and germline variant detection, RNA-Seq expression estimation and HLA typing.

This pipeline starts with raw patient tumor exome and cDNA capture(3) or RNA-seq data and produces annotated VCFs for neoantigen identification and prioritization with pVACtools. Our pipeline consists of several modular components: DNA alignment, HLA typing, germline and somatic variant detection, variant annotation, and RNA-seq analysis. More specifically, we use BWA-MEM(4) for aligning the patient's tumor and normal exome data. For germline variant calling, the output BAM then undergoes merging (merging of separate instrument data) (Samtools(5) Merge), query name sort (Picard SortSam), duplicate marking (Picard MarkDuplicates), position sorting (Picard SortSam), and base quality recalibration (GATK BaseRecalibrator). GATK's HaplotypeCaller(6) is used for germline variant calling and the output variants are annotated using VEP(7) and filtered for coding sequence variants.

For somatic variant calling, the pipeline is consistent with above except that the variant calling step combines the output of four variant detection algorithms- Mutect2(8), Strelka(9), Varscan(10,11) and Pindel(12). The combined variants are normalized using GATK's LeftAlignAndTrimVariants where the indels are left-aligned and common bases are trimmed. Vt(13) is used to split multi-allelic variants. Several filters such as gnomAD allele frequency (maximum population allele frequency), percentage of mapq0 reads, as well as pass-only variants are applied prior to annotation of the VCF using VEP. We use a combination of custom and standard plugins for VEP annotation (parameters: --format VCF --plugin Downstream --plugin Wildtype --symbol --term SO --transcript_version --tsl

--coding_only --flag_pick --hgvs). Variant coverage is assessed using bam-readcount (https://github.com/genome/bam-readcount) for both the tumor and normal DNA exome data and this information is also annotated into the VCF output using VCF-annotation-tools (http://vatools.org).

Our pipeline also generates a phased-VCF file by combining both the somatic and germline variants and running the sorted combined variants through GATK ReadBackedPhasing.

For RNA-seq data, the pipeline first trims adapter sequences using flexbar(14) and aligns the patient's tumor RNA-seq data using HISAT2(15). Two different methods, Stringtie(16) and Kallisto(17), are employed for evaluating both the transcript and gene expression values. Additionally, coverage support for variants in RNA-seq data is determined with bam-readcount. This information is added to the VCF using VCF-annotation-tools and serves as an input for neoantigen prioritization with pVACtools.

Optionally, our pipeline can also run HLA-typing *in silico* using OptiType(18) when clinical HLA typing is not available.

**Analysis of TCGA data using pVACtools**
To demonstrate the utility and performance of the pVACtools package, we downloaded exome sequencing and RNA-Seq data from The Cancer Genome Atlas (TCGA)(19) from 100 cases each of melanoma, hepatocellular carcinoma and lung squamous cell carcinoma, and used patient-specific MHC Class I alleles (**Supp Fig 1**) to determine neoantigen candidates for each patient. There were a total of 64,422 VEP-annotated variants reported across 300 samples, with an average of 214 variants per sample. Of these, 61,486 were single nucleotides variants (SNVs), 479 were inframe insertions and deletions and 2,465 were frameshift mutations (**Supp Fig 2**). We used this annotated list of variants as input to the pVACseq component of pVACtools to predict neoantigenic peptides. pVACseq reported 14,599,993 unfiltered neoantigen candidates. The original version of pVACseq(20) reported 10,284,467 neoantigens. This demonstrated that by extending support for additional variant types as well as prediction algorithms (due to support for additional alleles), we produced 42% more raw candidate neoantigens.

After applying our default median binding affinity cutoff of 500 nM across all eight MHC Class I prediction algorithms, there were 96,235 predicted strong binding neoantigens, derived from 34,552 somatic variants (32,788 missense SNVs, 1,603 frameshift variants and 131 in-frame indels). This set of strong binders was further reduced by filtering out mutant peptides with median predicted binding affinities (across all prediction algorithms) greater than that of the corresponding wildtype peptide (i.e. mutant/wildtype binding affinity fold change > 1), resulting in 70,628 neoantigens from 28,588 variants (26,880 SNVs, 1,583 frameshift and 125 in-frame indels).

This set was subsequently filtered by evaluation of exome sequencing data coverage and our recommended defaults, as follows. By applying the default criteria of variant allele fraction (VAF) cutoff of > 25% in tumor and < 2% in normal sample, with coverage levels of at least 10X tumor coverage and at least 5X normal coverage, 10,730 neoantigens from 4,891 associated variants (4,826 SNVs, 56 frameshift and 9 in-frame indels) were obtained, with an average of 36 neoantigens predicted per case. Since RNA-seq data also were available, the filtering criteria included RNA-based coverage filters (tumor RNA VAF > 25% and tumor RNA coverage > 10X) as well as a gene expression filter (FPKM > 1). To condense the results even further, only the top ranked neoantigen was selected per variant ('top-score filter') across all alleles, lengths, and registers (position of amino acid mutation within peptide sequence), resulting in 4,891 total neoantigens with an average of 16 neoantigens per case. This list was then processed with pVACvector to determine the optimum arrangement of the predicted high quality neoantigens for a DNA-vector based vaccine design (**Figure 3**).

**Comparison of filtering criteria**

Since pVACtools offers a multitude of ways to filter a list of predicted neoantigens, we evaluated and compared the effect of each filter for selecting high quality neoantigen candidates. There were 14,599,993 unfiltered neoantigen candidates predicted by pVACseq resulting from 64,422 VEP-annotated variants reported across 300 samples.

We first compared the effect of running pVACtools with the commonly used standard binding score cutoff of <= 500 nM (parameters: -b 500) to the newly added allele-specific score filter (parameters: -a). These cutoffs were, by default, applied to the "median" binding score of all prediction algorithms. In total, 96,235 neoantigens (average 320.8 per patient) were predicted using the 500 nm binding score cutoff compared to 94,068 neoantigens (average 313.6 per patient) using allele-specific filters. About 79% of neoantigens were shared between the two sets.

We also narrowed further to include only those predictions where the default "median" predicted binding affinities (<= 500 nm) are lower than each corresponding median wildtype peptide affinity (parameters: -b 500 -c 1) (i.e. a binding affinity mutant/wild type ratio or differential agretopicity value indicating that the mutant version of the peptide is a stronger binder) (21). Using the agretopicity value filter, 70,628 neoantigens (average 235.43 per patient) were predicted versus the previously reported 96,235 neoantigens without this filter.

We also evaluated the effect of the agretopicity value filter when applied to the set of peptides filtered on “lowest” binding score of <= 500 nm (parameters: -b 500 -c 1 -m lowest). This filter limits peptides to those where at least one of the algorithms predicts a strong binder, instead of calculating a median score and requiring that to meet the 500 nm threshold (**Figure 6**). Using the lowest binding score filter resulted in an 11-fold increase in the number of candidates predicted (827,423 candidates, average 2,758.08 per patient). This approach may be useful for finding candidates in tumors with low mutation burden (but may also presumably lead to a higher rate of false positives).

Using the median (default) binding affinity filtering criteria, we next applied coverage and expression based filters. First, we filtered using the recommended defaults i.e. greater than 5X normal DNA coverage, less than 2% normal VAF, greater than 10X tumor RNA and DNA coverage and greater than 25% tumor RNA and DNA VAF, along with FPKM > 1 for transcript level expression (parameters: --normal-cov 5 --tdna-cov 10 --trna-cov 10 --normal-vaf 0.02 --tdna-vaf 0.25 --trna-vaf 0.25 --expn-val 1). A total of 10,730 neoantigens were shortlisted across all samples with an average of 35 neoantigens per case. We then compared this set with a slightly more stringent criteria using tumor DNA and RNA VAF of 40% (parameters: --tdna-vaf 0.40 --trna-vaf 0.40). This shortened our list of predicted neoantigens to 4,073 candidates with an average of 13 candidates per patient.

**Demonstration of neoantigen analysis using pVACfuse**

To demonstrate the potential of neoantigens resulting from gene fusions, we analyzed TCGA prostate cancer RNA-seq data from 302 patients. This dataset was previously used as a demonstration set for the fusion neoantigen prediction supported by INTEGRATE-Neo(22). We wanted to assess the difference (if any) in neoantigens candidates reported by INTEGRATE-Neo using the one MHC Class I prediction algorithm it supports (NetMHC) versus an ensemble of eight Class I prediction algorithms supported in pVACfuse.

Using 1,619 gene fusions across 302 samples as input, pVACfuse reported 2,104 strong binding neoantigens (binding affinity <= 500nM) resulting from 739 gene fusions. On average, there were about 7 neoantigens per sample resulting from an average of 2 fusions per case. This is an eight fold increase in the number of strong binding neoantigens predicted by pVACfuse versus those reported by INTEGRATE-Neo alone, which reported 261 neoantigens across 210 fusions.

### Supplementary Figures

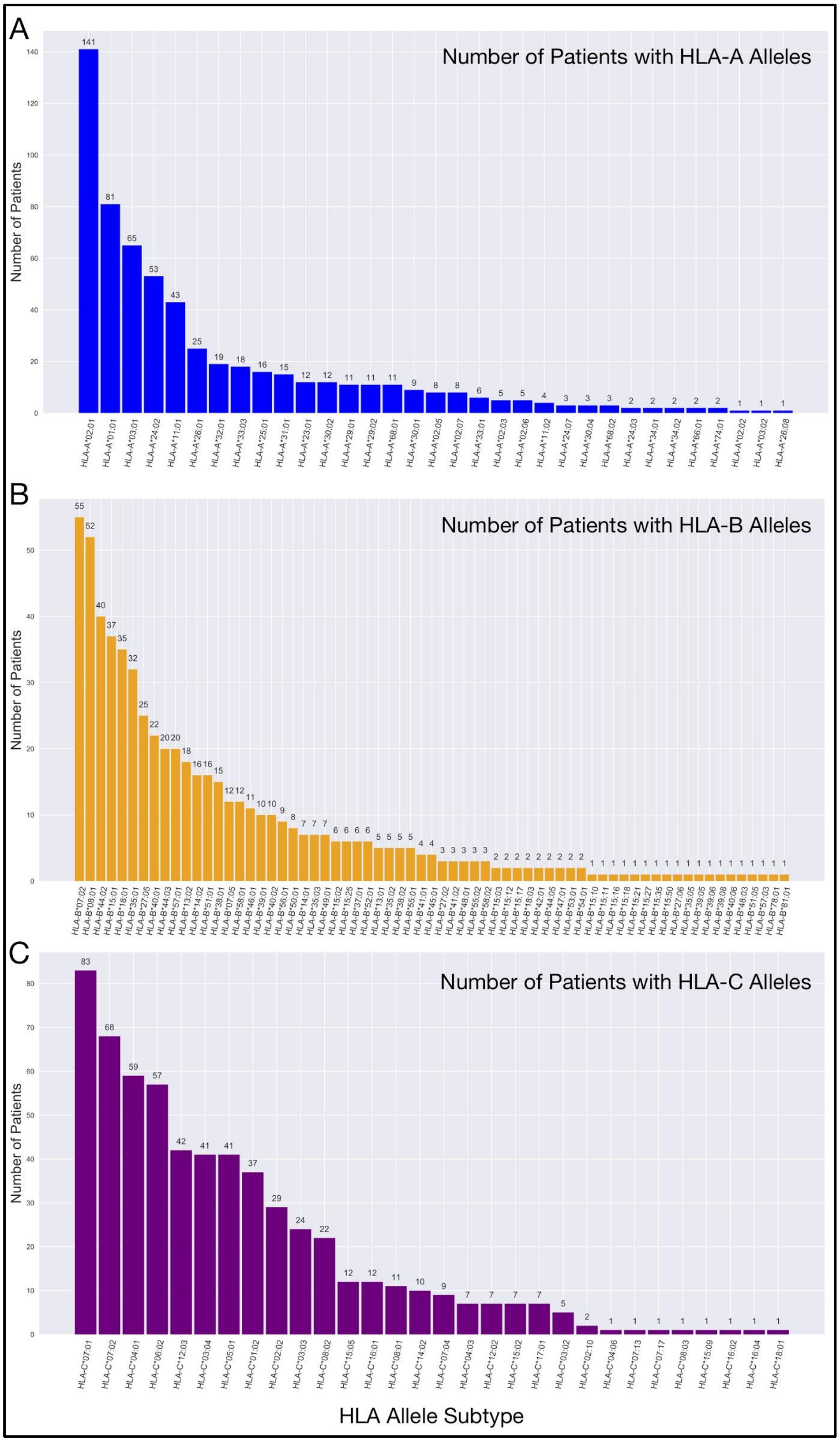


**Supp Fig 1: Patient counts per HLA allele subtype.** The distribution of HLA-alleles for the entire cohort of 300 TCGA patients analyzed in this study is shown in each of the three plots for (a) HLA-A allele subtypes (33 unique alleles observed); (b) HLA-B allele subtypes (67 unique alleles); and (c) HLA-C allele subtypes (30 unique alleles). The total number of unique Class I HLA alleles found in the patient cohort was 130.

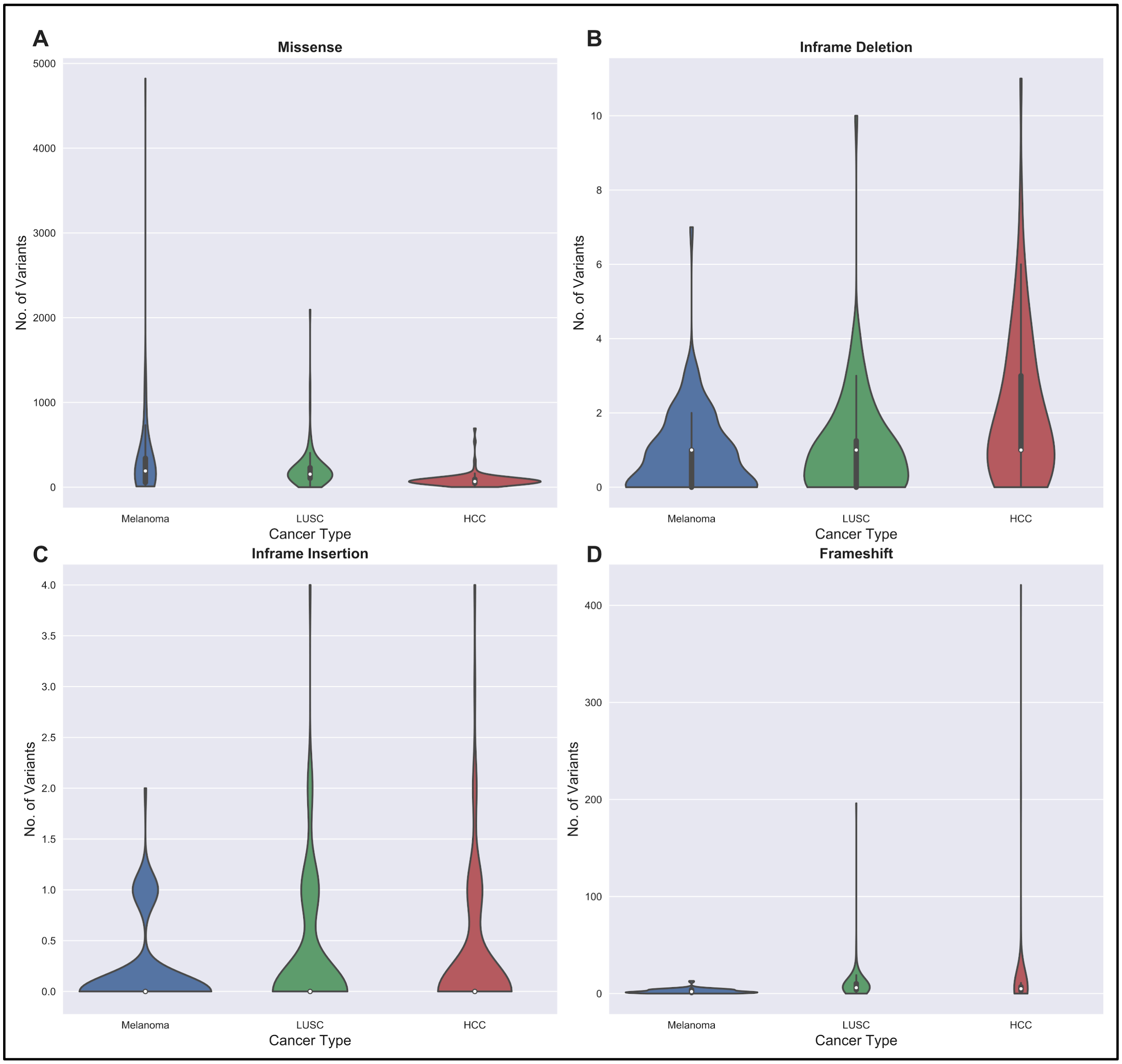


**Supp Fig 2: Number of variants per patient per cancer type** Violin plots show the distribution of observed variants per cancer type summarized for each variant type supported by pVACseq for (A) missense (total variant count: 61,486); (B) inframe deletion (total variant count: 389); (C) inframe insertion (total variant count: 81); and (D) frameshift (total variant count: 2,465).

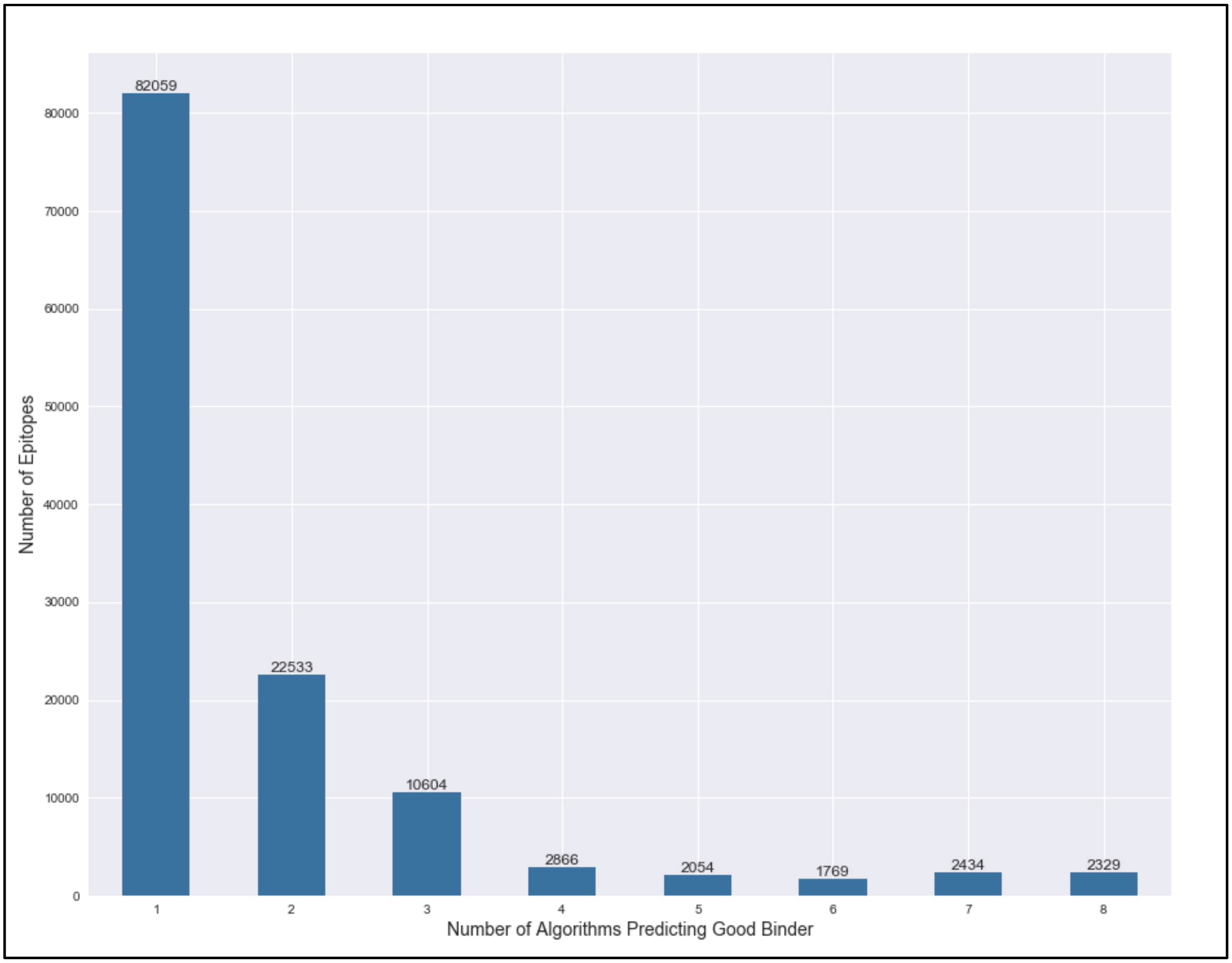


**Supp Fig 3:** Number of peptides predicted to be good binders (MT IC50 Score < 500 nM) versus number of algorithms used. Total number of 126,648 strong binding epitopes from 300 TCGA patients were analyzed.

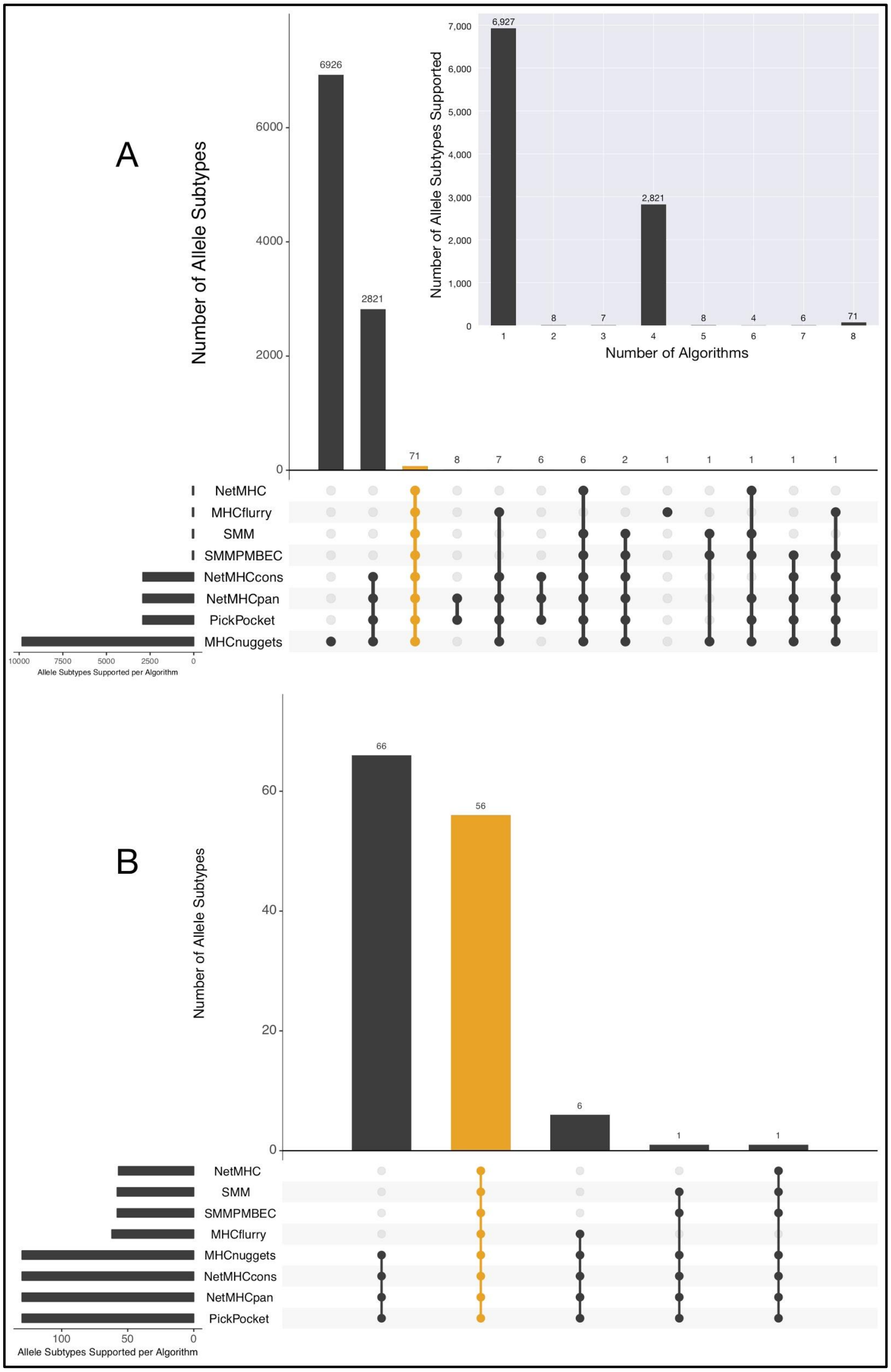


**Supp Fig 4:** Human HLA Allele subtype support for eight algorithms. (a) Number of allele subtypes supported versus number of algorithms; (b) Upset plot for algorithm combinations ranked by the number of allele subtypes supported by pVACseq. Total number of HLA allele subtypes supported by pVACseq: 9,851. The combination of all eight algorithms (highlighted orange) ranks the 3rd highest; (C) Upset plot for algorithm combinations ranked by the number of allele subtypes supported for the 300 TCGA samples analyzed in this study. Total number of HLA alleles across patient cohort: 130. The combination of all eight algorithms (highlighted orange) ranks the 2nd highest.

### Supplementary Tables

**Supp Table 1:** Comparison of existing software and tools for neoantigen detection and characterization

| | **pVACtools** | **Vaxrank**(23) | **MuPeXI**(24) | **CloudNeo** (25) | **FRED2**(26) | **Epi-Seq**(21) | **ProTECT** (27) |
|---|---|---|---|---|---|---|---|
| **Variant calling (within the pipeline)** | N | N | N | N | N | Y | Y |
| **Variant types supported** | SNVs, Inframe Indels, Frameshifts, Gene fusions (pVACfuse) | SNVs, Indels, | SNVs, Inframe Indels, Frameshifts | SNVs | SNVs, Inframe Indels, Frameshifts, | SNVs | SNVs |
| **HLA typing (within the pipeline)** | N | N | N | Y (PolySolver, HLAMiner) | Y (via separate installation of OptiType, Polysolver, Seq2HLA, & ATHLATES) | N | Y (Phlat) |
| **RNA-Seq expression data filter** | Y (pVACseq) | Y | Y | N | N | Y | Y |
| **Sequence coverage & VAF filter** | Y (pVACseq) | Y | Y (only compatible with MuTect2 variant calls) | N | N | Y | N |
| **Algorithms used for epitope binding affinity prediction** | IEDB (web and local)<br><br>(MHC Class I: NetMHCpan, NetMHC, NetMHCcons, PickPocket,, SMM, SMMPMBEC)<br><br>MHCflurry MHCnuggets<br><br>(MHC Class II: NetMHCIIpan, SMMalign, NNalign MHCnuggets) | IEDB (web and local)<br><br>MHCflurry, NetMHC, NetMHCpan, NetMHCIIpan, NetMHCcons, | netMHCpan (local only) | netMHC, netMHCpan (local only) | Local: NetMHC, NetMHCPan, NetMHCII, NetMHCIIpan, NetCTLpan, PickPocket<br><br>Included: Select from, SMMPMBEC, syfpeithi, SMM, Tepitopepan, ARB,epidemix, comblibsidney, Unitope, HAMMER, SVMHC, BIMAS, | NetMHC (local only) | IEDB (local only) |
| **Stability prediction** | Y (NetMHCstabpan) | N | N | N | Y (NetMHCstabpan) | N | N |

| | | | | | | | |
|---|---|---|---|---|---|---|---|
| **Cleavage site prediction** | Y (NetChop) | Y (NetChop) | N | N | Y (ProteaSMM, PCM, Ginodi, NetChop) | N | N |
| **Support for vector design/epitope assembly** | Y (pVACvector) | N | N | N | Y (OptiVac) | N | N |
| **Incorporation of proximal variants** | Y | Y (RNA only) | N | N | N | Y (RefHap) | N |
| **Unique epitope ranking method** | Y | Y (“Total Binding Score”) | Y (“Priority Score”) | N | Y (OptiTope immunogenicity score) | N | Y (Rankboost score) |
| **Graphical User Interface** | Y (pVACviz) | N | Y (http://www.cbs.dtu.dk/services/MuPeXI/) | Y (via NCI Cancer Genomics Cloud version) | Y (EpiToolKit 2.0) | N | N |
| **Results visualization** | Y (pVACviz) | N | N | N | N | N | N |
| **HTTP REST API** | Y (pVACapi) | N | N | N | Y | N | N |
| **License** | NPOSL-3.0 | Apache 2.0 | Unknown | Apache 2.0 | 3-clause BSD | Unknown | Apache 2.0 |

14. Roehr JT, Dieterich C, Reinert K. Flexbar 3.0 - SIMD and multicore parallelization.

Bioinformatics. 2017;33:2941–2.

27. ProTECT https://github.com/BD2KGenomics/protect
